# Hypothalamic C2-domain protein involved in MC4R trafficking and control of energy balance

**DOI:** 10.1101/793000

**Authors:** Chaitanya K Gavini, Tyler M Cook, David J Rademacher, Virginie Mansuy-Aubert

## Abstract

**Objective:** Rates of overweight and obesity epidemic have risen significantly in the past few decades, and 34% of adults and 15-20% of children and adolescents in the United States are now obese. Melanocortin receptor 4 (MC4R), contributes to appetite control in hypothalamic neurons and is a target for future anti-obesity treatments (such as setmelanotide) or novel drug development effort. Proper MC4R trafficking regulation in hypothalamic neurons is crucial for normal neural control of homeostasis and is altered in obesity and in presence of lipids. The mechanisms underlying altered MC4R trafficking in the context of obesity is still unclear. Here, we discovered that C2CD5 expressed in the hypothalamus is involved in the regulation of MC4R endocytosis. This study unmasked a novel trafficking protein nutritionally regulated in the hypothalamus providing a novel target for MC4R dependent pathways involved in bodyweight homeostasis and Obesity.

**Methods:** To evaluate the expression of C2cd5, we first used *in situ* hybridization and RNAscope technology in combination with electronic microscopy. For *in vivo*, we characterized the energy balance of wild type (WT) and C2CD5 whole-body knockout (C2CD5KO) mice fed normal chow (NC) and/or western-diet (high-fat/high-sucrose/cholesterol) (WD). To this end, we performed comprehensive longitudinal assessment of bodyweight, energy balance (food intake, energy expenditure, locomotor activity using TSE metabolic cages), and glucose homeostasis. In addition, we evaluated the consequence of loss of C2CD5 on feeding behavior changes normally induced by MC4R agonist (Melanotan, MTII) injection in the paraventricular hypothalamus (PVH). For *in vitro* approach, we tease out the role of C2CD5 and its calcium sensing domain C2 in MC4R trafficking. We focused on endocytosis of MC4R using an antibody feeding experiment (in a neuronal cell line - Neuro2A (N2A) stably expressing HA-MC4R-GFP; against HA-tag and analyzed by flux cytometry).

**Results:** We found that 1) the expression of hypothalamic C2CD5 is decreased in diet-induced obesity models compared to controls, 2) mice lacking C2CD5 exhibit an increase in food intake compared to WT mice, 3) C2CD5 interacts with endocytosis machinery in hypothalamus, 4) loss of functional C2CD5 (lacking C2 domain) blunts MC4R endocytosis *in vitro* and increases MC4R at the surface that fails to respond to MC4R ligand, and, 5) C2CD5KO mice exhibit decreased acute responses to MTII injection into the PVH.

**Conclusions:** Based on these, we conclude that hypothalamic C2CD5 is involved in MC4R endocytosis and regulate bodyweight homeostasis. These studies suggest that C2CD5 represents a new protein regulated by metabolic cues and involved in metabolic receptor endocytosis. C2CD5 represent a new target and pathway that could be targeted in Obesity.

## 1. Introduction

Obesity is a public health concern with over a 1.5 billion adults and children categorized as obese [1,2,3]. Within the United States, 33% of adults and 20% of children are estimated to be obese [4]. Based on the current trends and projections, increase in obesity prevalence is estimated to reach 60% in adult men, 40% in adult women, and 25% in children by 2050 [5]. Obesity results from a long-term positive energy balance, i.e., increased food intake and decreased energy expenditure [6]. Distinct neuronal circuits and signaling mechanisms within the brain, in particular, hypothalamus regulates energy balance (feeding and metabolism) through adjustments in physiological processes. Disruption of some of these hypothalamic mechanisms is thought to be involved in the obesity and had been targeted to improve energy balance homeostasis [7–12]. One such mechanism involves the melanocortin system. The ability of melanocortin to suppress feeding and increase energy expenditure has made melanocortin receptors (MCR) the focus of many anti-obesity treatments. Melanocortin system controls many aspects of metabolism that can contribute to obesity and associated comorbidities [13]. Studies on spontaneous as well as genetically introduced variations in genes of the melanocortin system demonstrate the importance of this system in obesity and its associated metabolic syndrome, and suggest a dysfunction of the melanocortin system [13,14].

One of the first melanocortin system genes studied using genetic deletion in mice was MC4R, a G-protein-coupled receptor. Disruption of MC4R in mice resulted in an obese phenotype, underscoring the potential role of MC4R in obesity [15,16]. Mutations in the MC4R are a relatively common cause of severe obesity [17,18,19,20]. Humans with MC4R mutations led to either gain [21] or loss of function [17,18]. Many of these mutations are suggested to cause either retention of the mutated receptors intracellularly, resulting in loss of agonist response [22,23], or altered internalization or recycling [21,24].

MC4R agonists lower food intake and BW [25]. Notably, recent Phase 3 trials of setmelanotide, a MC4R agonist, in obese patients have met their primary endpoints positioning it to bring the drug to market in 2020. Among the participants who hit a weight loss threshold after 12 weeks, the mean reduction in body weight over the course of the trial was 25.4%. It has been reported a 27.8% drop in hunger in those participants. Many current clinical trials aim at decreasing food intake and target MC4R signaling and trafficking. It is crucial to understand the neurobiology underlying the drug effects and to better define MC4R activation and cycling. In this study we identify a new MC4R trafficking protein that could be targeted in Obesity.

It has been known for some time that hypercaloric diets containing saturated fatty acid, induce insulin and leptin resistance, loss of MC4R ligand α-MSH) abundance, endoplasmic reticulum stress and inflammation [24]. Previous studies showed that during lipid stress MC4R internalization to endosomes as well as receptor routing to lysosomes is impaired leading to altered desensitization [24]. Specific proteins such as MRAP involved in MC4R trafficking to the plasma membrane had been extensively studied [26,27]. Human mutation that may affect MC4R endocytosis or recycling [21] had been recently shown to impact bodyweight homeostasis however, the specific protein(s) involved in MC4R endocytosis/recycling process are not identified yet.

Here, we identify a calcium- and lipid- binding C2 domain protein, C2CD5 [28] that contribute to regulation of MC4R endocytosis. C2CD5 is highly expressed in the hypothalamus and its level decreases in obesity and in presence of saturated fatty acids. Our data suggest that hypothalamic C2CD5 is a novel receptor trafficking protein that responds to metabolic cues and is involved in neural control of energy balance.

## 2. Materials and methods

### 2.1. Animal care and use

All studies were conducted in accordance to recommendations in the Guide for the Care and Use of Laboratory Animals of the National Institutes of Health and the approval of the Loyola University Chicago Institutional Animal Care and Use Committee. C57BL/6J (#000664) were obtained from Jackson laboratory (Maine, USA) and C2CD5KO were obtained from Texas A&M Institute for Genomic Medicine (College Station, Texas). Mice with identical genetic background were compared in all studies. Mice were genotyped and protein knockout was assessed using immunoblots (Supplementary 2). All mice were housed 4/cage under a 12:12 h light/dark cycle. Mice received either NC (Teklad LM-485) or WD (TD88137, Teklad Diets; 42%kcal from fat, 34% sucrose by weight, and 0.2% cholesterol total) (Envigo, Indiana, USA) for 12 weeks after initial assessment of energy balance and glucose homeostasis before BW divergence (~ week 7-8 of age). BW were recorded weekly from weaning. All metabolic studies mentioned were done using both male and female mice littermates with experimenter blinded to both treatment and genotype using the ARRIVE guidelines.

### 2.2. Cell culture

N2A cells stably transfected with HA-MC4R-GFP (kindly provided by Dr. Giulia Baldini) were cultured in DMEM with L-glutamine and sodium pyruvate supplemented with 10%FBS and penicillin/streptomycin. The N2A cells stably expressing HA-MC4R- GFP have been previously described [29].

### 2.3. Glucose and insulin tolerance tests

Glucose and insulin tolerance testing were performed as previously described [30,31,32]. For glucose tolerance, overnight (12hrs) fasted mice were given i.p dose of glucose (1g/kg BW) after measuring fasting glucose levels. Blood glucose levels were then monitored at 15, 30, 60, 120 min post injection using AlphaTrak glucometer for rodents (Fisher Scientific, Pennsylvania, USA). For insulin tolerance, mice were fasted for 4hrs and given i.p dose of insulin (0.5U/kg BW, Human-R Insulin U100, Lilly) with glucose levels monitored before and after as described above.

### 2.4. Western blotting

Hypothalamus or whole cell protein isolation and western blotting were performed as described before [30]. Briefly, whole cells or hypothalamus were isolated, homogenized in ice cold lysis buffer (1XPBS + 1% Triton X-100 + 0.01% SDS + protease/phosphatase inhibitor). Equal protein was loaded and resolved using 4–15% Mini-PROTEAN Precast Protein Gels (Bio-Rad, #4561086) and transferred onto PVDF membranes (Bio-Rad), and, processed for actin (abcam, #ab8226), HA (Biolegend, #901513), and, C2CD5 (#A301-469A, Bethyl laboratories, Montgomery, TX) at supplier recommended dilutions. Immunoblots were analyzed using ImageJ and normalized to actin levels and graphed using control groups as 100%.

### 2.5. Metabolic cage analysis

Metabolic measurements were performed using TSE Phenomaster (TSE systems, Chesterfield, MO) on WT and C2CD5KO mice before BW divergence to avoid confounding effects of BW on energy balance [33] and as described before [31]. Energy balance parameters were also assessed in the metabolic cages using a pair-feeding paradigm before BW diversion.

### 2.6. Immunoprecipitation

To identify proteins that interact with C2CD5, hypothalamus from 10-week-old male WT mice were used. Hypothalami were homogenized and protein levels determined (BCA protein assay kit, Thermo Fisher, # 23235). Five-hundred micrograms of protein was incubated with 1μg of rabbit polyclonal anti-C2CD5 antibody (Bethyl laboratories, #A301-469A), and rotated overnight at 4°C. Overnight samples were incubated with 25μl of protein A/G magnetic beads (Thermo Fisher, #88803) rotating for 4hrs at 4°C. Proteins were eluted from the beads with 50μl of lysis buffer containing β-mercaptoethanol and analyzed using mass spectrometry as described below.

To study the complex comprising C2CD5, AP2, and HA-MC4R-GFP in N2A cells, co-IP experimental conditions using anti-C2CD5 or anti-HA (1:150, Biolegend, #901513) were used as described above.

### 2.7. Proteomics and MS analysis

For C2CD5 IP’s, gel lanes/bands were excised, cleaned up, destained and subjected to in-gel tryptic digestion. The resulting peptides were run through LC-MS/MS. The raw data files were processed and quantified with searches performed against the UniProt database by using the PEAKS software with protein confidence score −10lgP more than 20 regarded as high confidence.

### 2.8. DNA Constructs

A full-length cDNA clone of human *KIAA0528* (accession #BC117143) was obtained from Dharmacon Research, Inc with IMAGE clone ID 40125694. The clone was provided in a pCR4-TOPO vector. The mutants of C2CD5-mCherry, and ΔC2, were made by the PCR-based site-directed mutagenesis are described below and the DNA sequences for all expression constructs were confirmed before using in the expression studies and were validated for protein expression using immunoblotting. To make C2CD5-ΔC2 expression vector, the first 405 nucleotides were removed from the 5’ end of C2CD5 cDNA using Mlu1 (MCS site) and Cla1 (405) restriction enzymes and a polylinker was used to ligate the remaining cDNA in right flame to the pCMV5 vector. To express full-length C2CD5 fused with mCherry at its C-terminus, the stop codon before the BamH1 site was removed using PCR-based mutagenesis, and the cDNA ligated into the pcDNA3-mCherry vector at the Mlu1 and BamHI sites.

### 2.9. Stereotaxic surgery

Stereotaxic surgeries were performed to chronically implant guide cannulae (Plastics One Inc., Roanoke, VA, USA) aimed at the PVH. Mice were anaesthetized using isoflurane and mounted on a stereotaxic apparatus with atraumatic ear bars (Stoelting Co., Wood Dale, Illinois). The following coordinates obtained from Paxinos, George & Franklin, Keith BJ (Mouse brain in stereotaxic coordinates, 2001) were used for the PVH: anterior–posterior, −0.55mm; medial–lateral, −0.2mm; dorsal–ventral, −4.6mm and an injection needle with 0mm projection.

### 2.10. Melanotan II (MTII) treatment

After recovery from stereotaxic surgery, metabolic parameters were measured in WT and C2CD5KO mice. Mice were microinjected with either vehicle, aCSF (Harvard Apparatus, #59-7316) or MCR agonist, MTII (30pmoles/200nL) (Phoenix pharmaceuticals, Virginia, USA, # 043-29) over 30sec; the needle was left in place for an additional 10–15sec to minimize potential flow up the cannula track. The impact of MCR agonist on food intake was also assessed. For this, mice were deprived of food overnight (~16hrs) before being microinjected with MTII or aCSF, in order to detect a decrease in food intake. After the microinjections, mice were placed back into the metabolic cages. Food intake was determined at 2, 4, 6, 12, 24, and 48hrs post injection as described above. BW measurements were also taken. The study was repeated after a 4-day period such that each mice was injected with aCSF and MTII (order counterbalanced), to allow sufficient washout time to prevent any residual drug effects. Any possible variability in feeding efficiency due to minor differences in BW within group is accounted for by the repeated-measures design and by measuring both aCSF and MTII effects within the same mice.

### 2.11. In situ hybridization

Fluorescence in situ hybridization (FISH) was performed on 20 μm thick slices of fresh frozen mouse brains using RNAscope fluorescent multiplex reagents (Advanced Cell Diagnostics, 320850) according to the manufacturer’s instructions. RNA probes for C2cd5 and Mc4r (Advanced Cell Diagnostics, 436771 and 319181-C3 respectively) were incubated with the brain slices and signal amplification was achieved using the multiplex reagents as instructed. Images were captured using Olympus IX80 Inverted Microscope equipped with an X-Cite 120Q fluorescent light source (Lumen Dynamics) and a CoolSNAP HQ2 CD camera (Photometrics). Image processing and RNA signal quantification was done using CellSens software (Olympus Corporation, Waltham, Massachusetts).

### 2.12. Determination of HA-MC4R-GFP at the cell surface by flow cytometry

Total MC4R present at the cell surface was determined by flow cytometry using LSRFortessa (BD Biosciences, San Jose, CA). Briefly, N2A (HA-MC4R-GFP) cells were transfected with either C2CD5-mCherry or ΔC2-C2CD5-mCherry using Continuum transfection reagent (Gemini Bio, #400-700). Transfected cells were treated with FFA-free BSA or palmitate (200μM) for 18hrs. Live cells were washed with complete media and were incubated with anti-HA antibody (1:500, Biolegend, #901513) for 2hrs at 4°C prior to fixation. The HA-MC4R-GFP present at the cell surface was assessed using Alexa 647 conjugated antibody in non-permeabilized conditions. Total cell HA-MC4R-GFP was quantified by measuring the GFP fluorescence intensity. To determine the fraction of total receptor residing at the cell surface, the ratio HA-MC4R-GFP at the cell surface (Alexa 647 fluorescence)/total receptor (GFP fluorescence) was quantified for each condition.

To study the effect of α-MSH on the amount of MC4R at the cell surface, N2A cells from above groups were incubated for 1hr at 37°C with DMEM containing 100μM cycloheximide to inhibit protein synthesis. The cells were then incubated in the continuous presence of cycloheximide with or without 100 nM α-MSH for 30, 60, and 120min. Cells were fixed, and the cell surface MC4R was measured as described above.

### 2.13. cAMP assay

N2A cells from conditions mentioned above were treated with DMEM containing 0.5mM IBMX for 10min at 37°C, and then stimulated with 0.5mM IBMX and 200nM MTII for 15min at 37°C. Samples were collected in 0.1M HCl containing 0.5mM IBMX, centrifuged, and analyzed according to the manufacturer’s instructions (Enzo Life Sciences). The collected supernatants in 0.1M HCl were also used to determine protein concentration with the BCA protein assay.

### 2.14. ImmunoEM

Procedures to prepare brain tissue for electron microscopy were modified from Figge et al., 2013 and Yi et al., 2001 [34,35]. Briefly, WT mice were deeply anesthetized with isoflurane and transcardially perfused with 10 ml of PBS followed by 100 ml of PB containing 2% paraformaldehyde (Electron Microscopy Sciences, Hatfield, PA), 2.5% glutaraldehyde (Electron Microscopy Sciences), and 0.01% calcium chloride and coronal sections through the rostro-caudal extent of the hypothalamus were obtained. Images were acquired with a Philips CM120 transmission electron microscope with (TSS Microscopy, Beaverton, OR) equipped with a BioSprint 16 megapixel digital camera (Advanced Microscopy Techniques, Woburn, MA).

### 2.15. TIRF microscopy

For cell imaging, an Apo TIRF 60 × 1.49 numerical aperture oil-immersion objective with a penetration depth of around 100nm was used. N2A cells were transfected with either plasmid containing mCherry only or ΔC2-C2CD5-mCherry as described above in 35mm glass-bottom dishes (MatTak Corp, Ashland, Massachusetts). Positively transfected N2A cells were identified, and MC4R trafficking to the TIRF zone (GFP puncta) were recorded every 5min for a 30min period using Nikon Eclipse-Ti TIRF microscope and incubation system maintained at 37°C.

### 2.16. Quantification and statistical analysis

All data are represented as Mean±S.E.M. Analysis were done using IBM SPSS Statistics 24. For single group comparisons, either a 1- or 2-tailed t-test was used as appropriate and multiple comparisons were performed using ANOVA. For repeated measures, 2-way ANOVA was used and p value less than 0.05 was considered significant.

## 3. Results and Discussion

### 3.1. Expression of C2CD5 in the brain

Previously, C2CD5 has been shown to be involved in GLUT4 trafficking in adipocytes [28], and, in insulin sensitivity and browning of adipose tissue [36]. So far, the expression and function of C2CD5 in the brain has not been reported. In the present study, we show for the first time the presence of C2cd5 mRNA in the brain using *in situ* hybridization (Figure 1A). Results from in situ show marked expression throughout the brain with high presence in hypothalamus (Figure 1A1). We observed that mRNA and protein levels of C2CD5 are decreased in the hypothalamus of western diet-induced obesity model and *ex* vivo organotypic hypothalamic cultures treated with saturated fatty acids, (Figure 1B-E). These data suggested a role of C2CD5 in hypothalamic centers. We performed a high-resolution immuno-electron microscopy to visualize C2CD5 cellular localization in the hypothalamus. Immunogold labelling of hypothalamic tissue showed the presence of C2CD5 near the plasma membrane and with mitochondrial membrane (Figure 1F).

**Figure 1:**
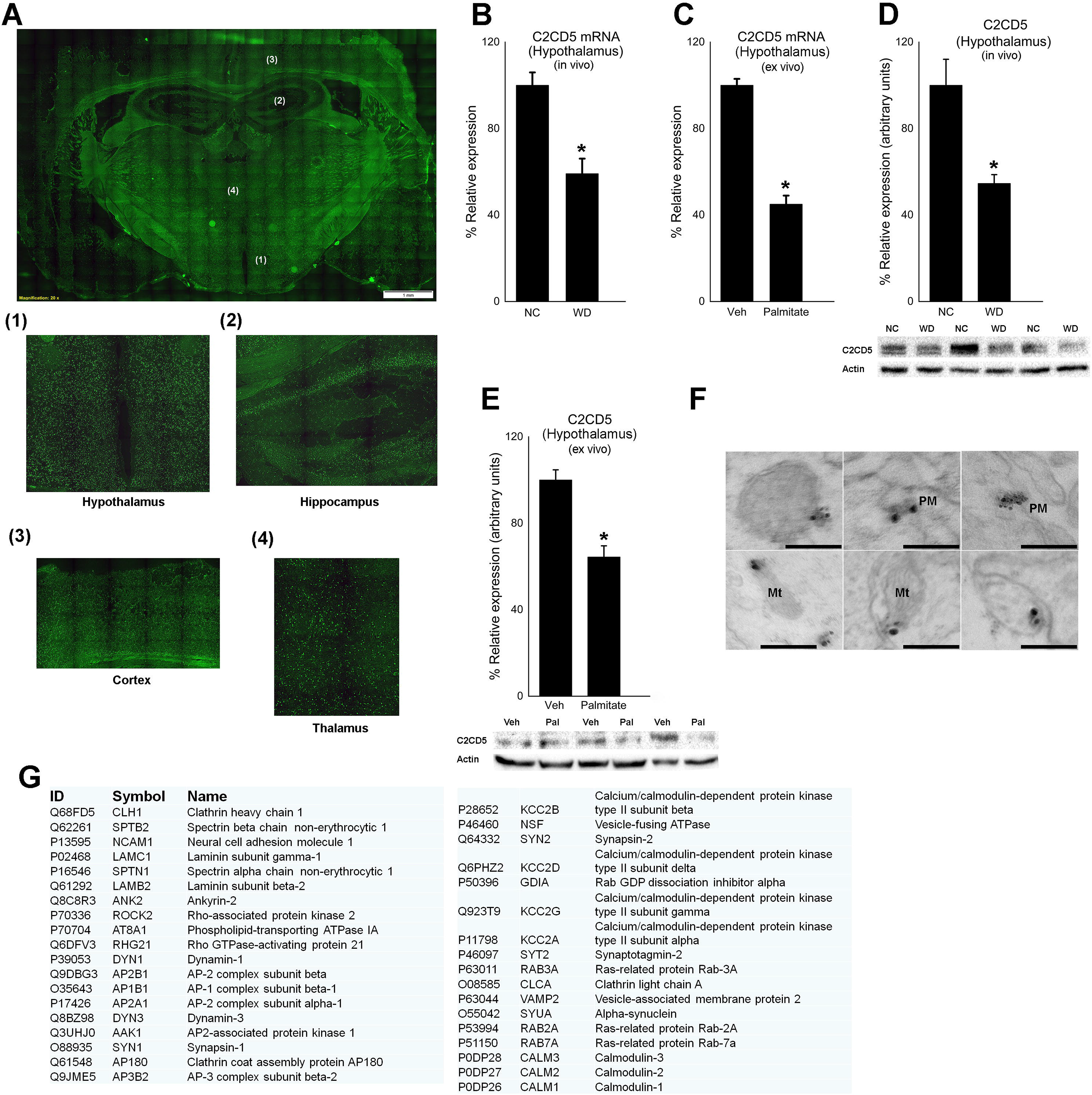
C2CD5 expressed in the brain interacts with endocytosis machinery and is downregulated in diet-induced obesity. A) mRNA expression of C2cd5 in the brain using in situ hybridization (Green, C2CD5 mRNA; Blue, Dapi). B, C) mRNA expression of C2cd5 in the hypothalamus of diet-induced obese mice (n=6/group) and hypothalamic organotypic cultures treated with 200μM palmitate (n=3 experiments in triplicates) respectively. D, E) Protein expression of C2CD5 in the hypothalamus of diet-induced obese mice (n=5/group) and hypothalamic organotypic cultures treated with 200μM palmitate (n=3 experiments in triplicates) respectively. F) Immunogold staining of mice hypothalamus for C2CD5 (PM, Plasma membrane; Mt, mitochondria; scale 200nm). G) Protein analysis of hypothalamic C2CD5 immunoprecipitation. C2CD5 interacts with components of plasma membrane and endocytosis machinery. All data are Mean±SEM. *p<0.05.

To check and ascertain the role of C2CD5 in the hypothalamus we looked for the interacting partners of C2CD5 in mice hypothalamus using immunoprecipitation followed by mass spectrometry (Figure 1G/ Supplementary Excel). Analysis from pull down of C2CD5 showed exclusively proteins involved in endocytosis machinery (e.g. AP2, clathrin heavy/light chains, dynamin1, 3) and components of plasma membrane. Based on these complementary approaches, we propose that C2CD5 is a novel hypothalamic protein responding to fat challenge, and is involved in trafficking in line with its role in adipose tissue [28].

### 3.2. Mice lacking C2CD5 develop obesity

There is increasing evidence that trafficking proteins could modulate the activity and cell surface expression of receptors, thereby regulating energy balance [22,26,27,37]. To determine the role of metabolically regulated trafficking protein C2CD5 in BW homeostasis, we compared energy balance parameters between C2CD5 whole body knockout (C2CD5KO homozygous) and WT controls. When fed a NC, male C2CD5KO mice were significantly heavier starting at 12 weeks of age compared to WT mice (Figure 2A), whereas female C2CD5KO had comparable BW to WT mice (Supplementary 1A). When fed a WD, male and female C2CD5KO mice were heavier than their NC-fed littermates respectively (Figure 2A, Supplementary 1A). Compared to WT mice, male C2CD5KO mice had higher blood glucose levels than WT mice at 18 weeks of age irrespective of diet (Figure 2B) (WT-NC < KO-NC < KO-WD). They also developed impaired glucose tolerance (Figure 2B) and insulin tolerance (Figure 2C). Both male and female C2CD5KO had elevated levels of circulating insulin, leptin, triglycerides, cholesterol, and increased lipid accumulation in the liver (Figure 2D-H, Supplementary 1D-G). These results indicate that male and female C2CD5KO mice develop obesity and associated metabolic syndrome when fed WD. However, NC-fed female mice lacking C2CD5 are similar to WT mice. This discrepancy between male and female is interesting to highlight because the pathology of obesity has a sexual dimorphism component with crucial role of estrogen and estrogen receptor [38]. In this study, we observed that only WD-fed female C2CD5KO differs from the WT mice. It is possible that a gonadal hormone such as estrogen compensates the lack of C2CD5 expression when mice are NC-fed but this compensation may be lost after chronic high-fat diet. Recent data shows the evidence of a sexual dimorphic nature of the brain in its response to high fat diet and in many cases, the use of chronic high fat diet exacerbates sexually dimorphic mouse obesity. Hubbard et al. showed that hypothalamic MC4R ligands are required for observing sexual dimorphism [39]. More investigations will be necessary to evaluate whether gonadal hormones regulate hypothalamic C2CD5 and define such a sexual dimorphic mouse obesity.

**Figure 2:**
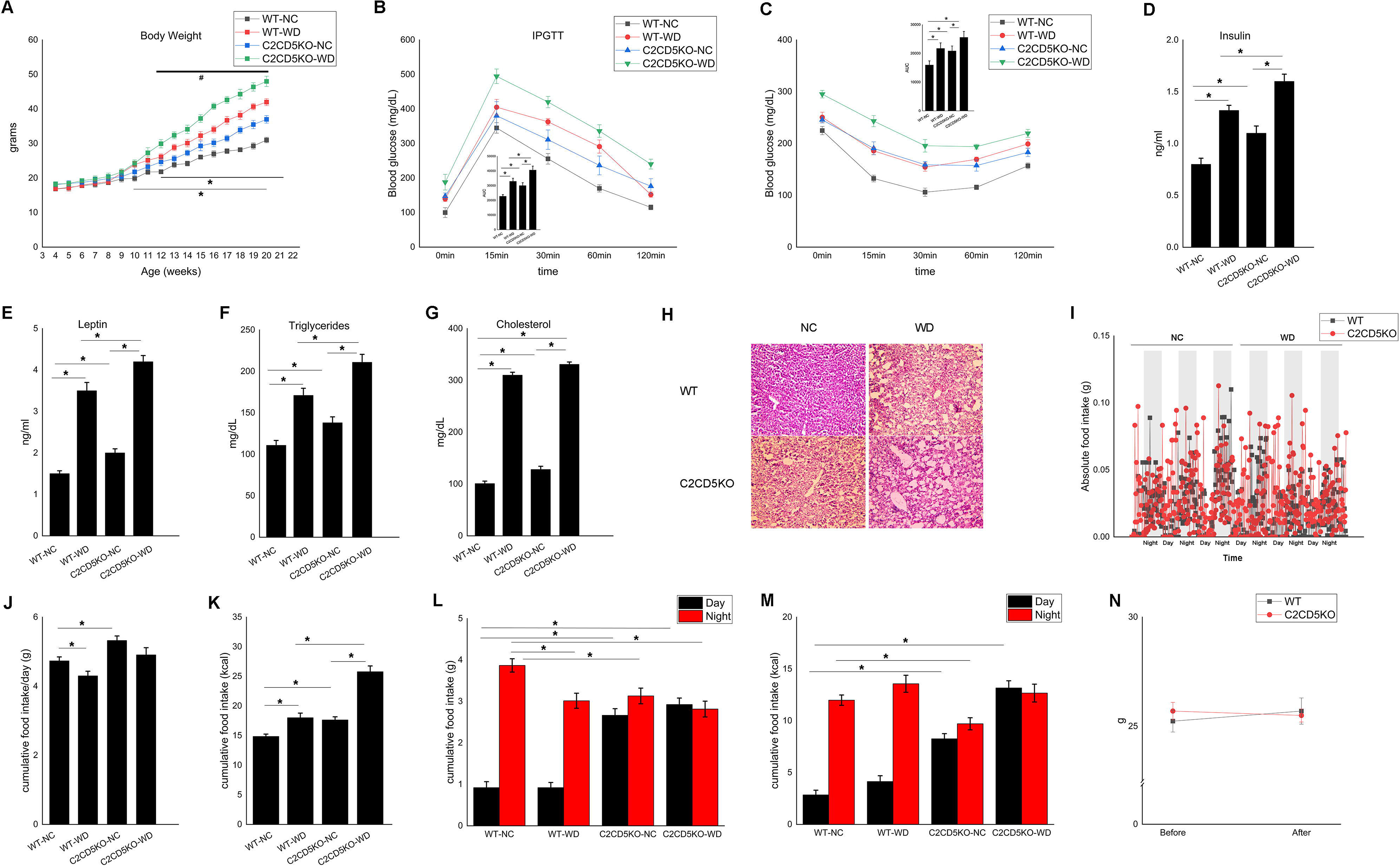
Loss of whole body C2CD5 results in obesity partially due to increased food intake. A) Body weight of male WT and C2CD5KO mice fed NC and WD. Mice were fed WD starting week10 of age just before body weights start to diverge (n=8-10/group) (* WT-NC vs WT-WD; # WT-WD vs C2CD5KO-WD; $WT-NC vs C2CD5KO-NC) B) Intra-peritoneal glucose tolerance test in male WT and C2CD5KO after 10 weeks of diet change (n=8/group). C) Intra-peritoneal insulin tolerance test in male WT and C2CD5KO after 10 weeks of diet change (n=8/group). D) Serum insulin levels of male WT and C2CD5KO after 12 weeks of diet change (n=6/group). E) Serum leptin levels of male WT and C2CD5KO after 12 weeks of diet change respectively (n=7/group). F,G) Serum triglycerides and cholesterol levels of male WT and C2CD5KO after 12 weeks of diet change respectively (n=8/group). H) Representative images of lipid accumulation in liver of WT and C2CD5KO when fed NC and WD. I) Absolute food intake of male WT and C2CD5KO mice before BW diversion when fed NC and WD respectively (n=8/group). J-M) Cumulative food intake in grams and kcal of male WT and C2CD5KO mice before BW diversion when fed NC and WD respectively (total and day/night) (n=8/group). N) BW of male WT and C2CD5KO mice during pair-feed paradigm (n=4/group). All data are Mean±SEM. *p<0.05.

### 3.3. Hyperphagia and reduced energy expenditure in C2CD5KO mice

Using metabolic cages, we assessed whether increased food intake or energy expenditure was a cause for increased BW and obesity in C2CD5KO mice. Before BW diversion, male C2CD5KO mice consumed ~20% and ~35% more calories than WT mice (8 weeks of age) when fed NC and WD respectively (Figure 2I-K). Male C2CD5KO mice significantly consumed more food during light phase with no significant differences seen during dark phase (Figure 2M). Before BW diversion, female C2CD5KO mice consumed significantly more calories than WT mice (8 weeks of age) only when fed WD (Supplementary 1H) consistent with the BW data (showing no difference between WT and KO when fed NC) and with the hypothesis that hypothalamic C2CD5 may be involved in the exacerbation of sexually dimorphic obesity induced by chronic western-diet [39]. Thus, lack of C2CD5 causes hyperphagia in males when fed NC or WD and in females when fed WD.

Energy expenditure was assessed using indirect calorimetry. In all groups, there was no significant difference in BW between groups avoiding any confounding effects of BW on energy balance [33]. As shown in Supplementary 2A, compared to WT mice, male C2CD5KO mice always exhibited significantly lower energy expenditure. However, female mice lacking C2CD5 did not show any significant differences in energy expenditure compared to WT mice (Supplementary 2B), suggesting that the hyperphagia observed in WD-fed female C2CD5KO mice drove obesity. Compared to WT mice, male C2CD5KO mice had a significantly decreased total activity and mostly during the dark cycle (Supplementary 2C), which might also contribute to reduced energy expenditure in these mice. No significant differences were observed in activity of the female mice (Supplementary 2D). The lack of energy expenditure phenotype in female cohort compared to male is again suggesting a sexual dimorphic role of hypothalamic C2CD5. Indeed, previous reports showed that estrogens through ERα regulate both the gene and protein expression (such as BDNF) in the hypothalamus which has been associated to neural modulation of white fat to induce browning [40,41]. More experiments using Cre lines (e.g hypothalamic nuclei and adipose specific), estrogen treatment and ovariectomized animals will define whether hypothalamic C2CD5 is involved in obesity related sexual dimorphism.

To determine the contribution of food intake and energy expenditure to obesity development in male C2CD5KO mice, we conducted a pair-feeding experiment before BW diversion. Male C2CD5KO mice were pair-fed in metabolic cages to restrict their daily food intake to the level of WT mice. Pair feeding did curtail the BW increase in male C2CD5KO mice (Figure 2N). Compared to WT, male C2CD5KO mice have a decreased energy expenditure (Supplementary 2E), this was masked (p<0.058) by slight increase in physical activity during pair feeding probably due to increased food anticipatory behaviors prior to feeding and spent more time eating after food became available [42]. Hence, increased food intake and a reduction in energy expenditure were in part responsible for the development of obesity in male C2CD5KO. These data from automated metabolic cages showed that C2CD5KO mice exhibited a robust increase in food intake before BW diversion.

### 3.4. C2CD5 regulates MC4R endocytosis and activity

As described above, the hypothalamus is a key regulator of food intake and BW, and the paraventricular hypothalamus (PVH) takes a crucial lead of this hypothalamic control of energy balance [43]. Studies have shown MC4R neurons of the PVH in the control of energy intake [44,45,46]. To determine if C2CD5 is expressed in the PVH and may modulate neuronal function, we examined the expression of C2cd5 in the PVH and in Mc4r expressing neurons using *in situ* hybridization. As shown in Figure 3A, C2cd5 is expressed in the PVH and we observed that 100% of Mc4r positive cells have C2cd5 mRNA.

**Figure 3:**
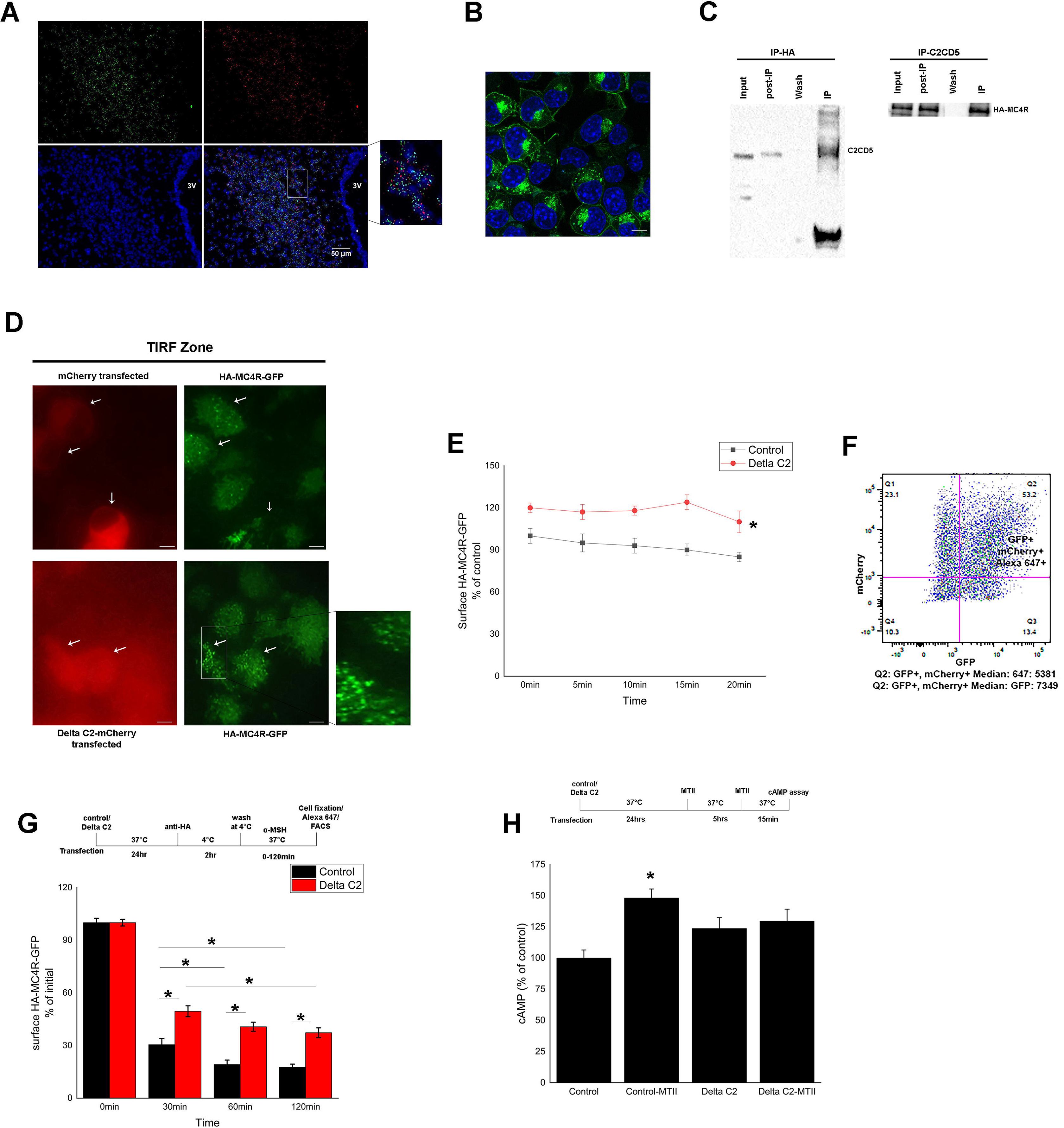
C2CD5 colocalize with MC4R in the PVH and regulates MC4R endocytosis. A) Colocalization of C2cd5 and Mc4r mRNA in the PVH (green, C2CD5; red, MC4R; blue, dapi; 3V, third ventricle; scale 50μm). B) Representative image of N2A cell line with stable expression of HA-MC4R-GFP (green, MC4R protein; blue, dapi; scale 10μm). C) C2CD5 interacts and co-precipitate with MC4R in N2A cells. D) MC4R at cell surface measured using TIRF microscopy in N2A cells transfected with mCherry or Delta C2-mCherry (red, mCherry or Delta C2-mCherry transfected cells; green, GFP puncta representing HA-MC4R-GFP at cell surface in TIRF zone; scale 10μm). E) Levels of MC4R at cell surface measured using GFP intensity in TIRF zone. F) Scatter plot of fluorescence intensity of GFP, mCherry, and Alexa 647 in N2A cells transfected with WT-C2CD5-mCherry or Delta C2-mCherry showing the feasibility of using flow cytometry for cell surface receptor quantification. G) α-MSH dependent decrease of MC4R at the cell surface in control or Delta C2 transfected N2A cells treated (n=4 experiments in triplicates). H) MTII-induced cAMP levels in N2A cells transfected with and without Delta C2, and treated with MTII for 5hr (n=3 experiments in triplicates). All data are Mean±SEM. *p<0.05.

Based on previous findings and our current work, C2CD5 is a trafficking protein [28] and loss of C2CD5 leads to increased food intake. Upon this, we hypothesized that role of C2CD5 in energy balance (e.g food intake and energy expenditure) could be the consequence of its role in MC4R trafficking or signaling. To study biochemical interactions between C2CD5 and MC4R, as there is no availability of a reliable antibody that can specifically target MC4R, we used a neuroblastoma cell line, Neuro2A (N2A) stably expressing HA-MC4R- GFP and extensively used in previous MC4R trafficking studies [24,29,47]. Co-IP experiments showed that C2CD5 and MC4R interact (Figure 3C).

As previously reported, C2-domain of C2CD5 is required to traffic GLUT4 in adipocytes [28]. We generated a mutant lacking the C2-domain (Delta C2 mutant) tagged with mCherry. Using TIRF microscopy (that images fluorescence of HA-MC4R-GFP in the TIRF zone representing up 70nM from the plasma membrane) we showed that the levels of MC4R near the plasma membrane was increased in the presence of Delta C2 mutant (Figure 3D, E) compared to controls. To better understand the underlying cause of the MC4R accumulation at the plasma membrane we used trafficking assay (to evaluate either endocytosis (or internalization) or exocytosis (or recycling)). First, we used antibody-feeding approach [29] followed by flow cytometry to evaluate α-MSH induced internalization. The rate of MC4R internalization was not significantly different between control cells and cell transfected with Wt-C2CD5 (not shown), whereas cells transfected with Delta C2 had a lower rate of receptor internalization (Figure 3F, G). This suggests that C2CD5 affects α-MSH induced internalization and that calcium and membrane binding is potentially involved because the lack of C2 domain decreases the internalization. We did not observe an effect of C2CD5 on recycling nor exocytosis (not shown). Then, we assessed whether the increase in MC4R receptor number at the surface (observed in cells lacking C2CD5) alter MC4R signaling. To this end, MTII was added for 5 hours to the medium of cells to induce MC4R signaling and cycling events (internalization, recycling, as previously shown [24]), in presence of functional C2CD5 or not. To test the capability of surface MC4R to generate cAMP, we treated cells with MTII again and measured cAMP after 15 min stimulation [24]. As shown in figure 3H, while the level of receptors at the surface is roughly 30% higher in cells with non-functional C2CD5, the cAMP level induced by MTII is not increased compared to control. These results suggest that while the loss of C2CD5 induces an increase in MC4R at the surface, these receptors are not functional potentially because they did not undergo proper recycling after being activated as suggested by Cooney et al., and Lotta et al. [21, 24]. It also suggests that gain of function mutation observed in human likely improves recycling but does not impair internalization only [21].

Previous work using N2A cell line showed that exposure to elevated levels of palmitate induced accumulation of MC4R at the surface but a decrease in the cAMP content to acute stimulation by α-MSH [48] suggesting non-functional receptors [24]. c2cd5 mRNA and protein expression is decreased in N2A cells treated with palmitate for 16-18hrs (Figure 4A, B). Our results with Delta C2 mutant mimic palmitate mediated changes in MC4R endocytosis levels and could be due to the decrease in C2CD5 expression. We hypothesized that expression of C2CD5 could rescue the palmitate-induced dysfunction in MC4R internalization. Among cells exposed to elevated levels of palmitate, cells transfected with Wt-C2CD5 had a better rate of MC4R internalization compared to control cells (Figure 4C, D). Consistent with previous report [24], N2A cells treated with palmitate (200μM) exhibit a blunted increase of cAMP following MTII acute exposure (~15 min) compared to vehicle (Figure 4E). To assess if the improvement of MC4R internalization in palmitate treated C2CD5 overexpressing cells has any functional relevance, we exposed Wt-C2CD5 overexpressing N2A cells to palmitate and treated with MTII. Palmitate-induced loss of the cAMP signal in the control N2A cells was rescued by the overexpression of Wt-C2CD5 (Figure 4E). MC4R, upon ligand binding, interacts with downstream proteins to direct signaling and this is attenuated by interacting with β-arrestins. This also promotes internalization of ligand bound receptors to early endosomes, from where it is either recycled to the cell membrane or moved to lysosomes for degradation [21]. Studies of others and our current suggest that treating cells with palmitate reduced the internalization of MC4R by limiting its interaction with endosomal machinery, thus suggesting exposure to elevated palmitate may impair internalization of MC4R and/or routing of the receptor to proper endosome/lysosomes [29,47,49]. Notably, palmitate [24] or lack of functional C2CD5 increase the MC4R at the surface by decreasing the internalization and proper recycling of MC4R. Given, the phenotype of C2CD5KO mice and diet-induced obesity models, these events likely lead to loss of function. In some cases, impaired internalization (or improved recycling) could also lead to gain of function if the level of receptor that increases at the surface are functional and not desensitized as suggested by Lotta et al., [21], however more investigation are needed to clearly linked MC4R trafficking events to metabolic phenotype.

**Figure 4:**
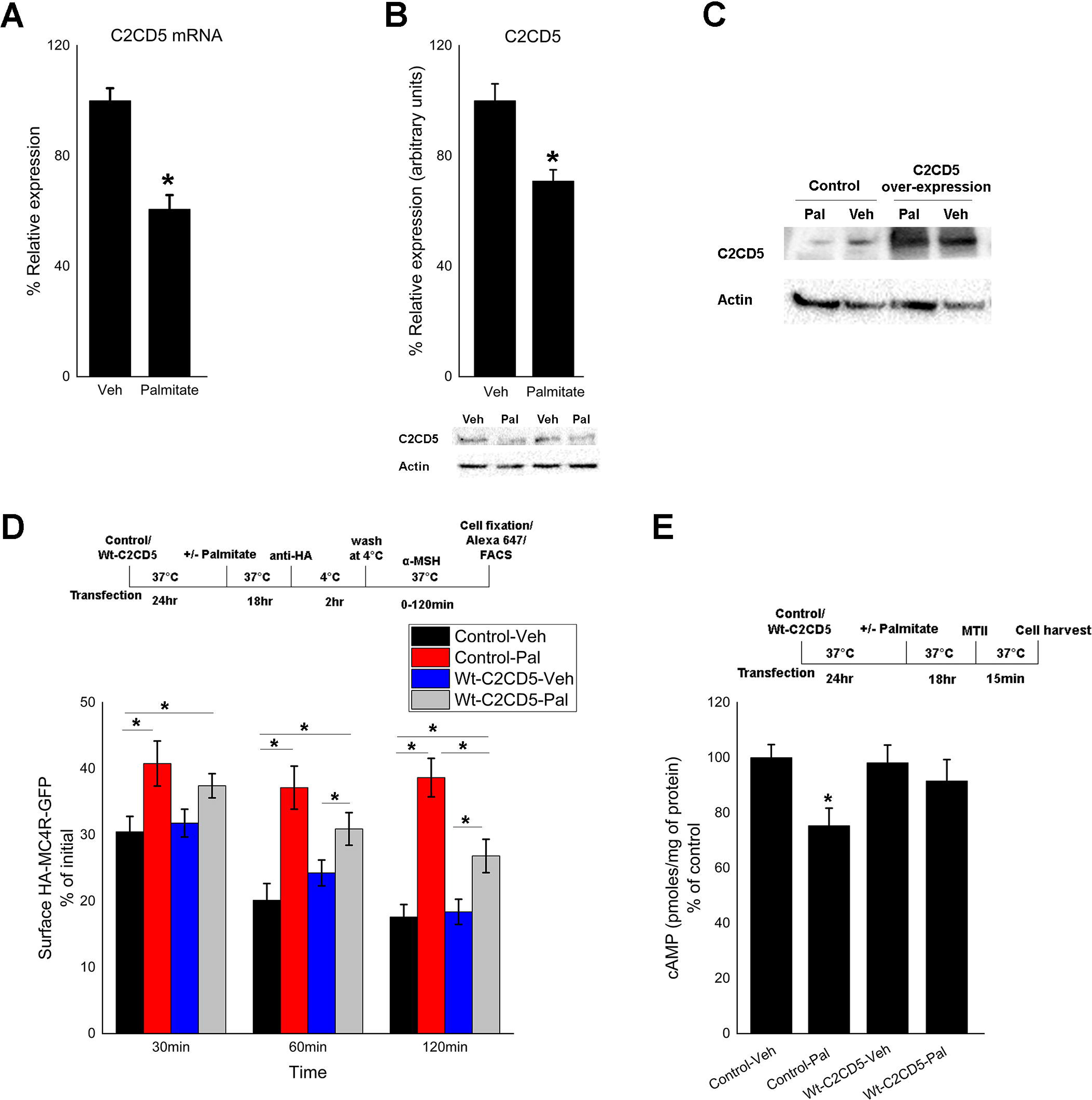
A, B) mRNA and protein expression of C2CD5 in N2A cells treated with vehicle or palmitate (200μM) respectively (n=3 experiments in triplicates). C) Western blot image showing overexpression of C2CD5 in Wt-C2CD5 transfected N2A cells. D)α-MSH dependent decrease of MC4R at the cell surface in control or Wt-C2CD5 transfected N2A cells treated with palmitate (n=4 experiments in triplicates). E) MTII-induced cAMP levels in control or Wt-C2CD5 transfected N2A cells treated with vehicle or palmitate (200μM) (n=3 experiments in triplicates). All data are Mean±SEM. *p<0.05.

### 3.5. Effects of MTII are impaired in mice lacking C2CD5

As discussed above, MC4R in the PVH regulate energy balance [44,45,46,50,51]. Disruption of MC4R activity in mice resulted in obese phenotype and increased axial length [15], underscoring the potential role of MC4R in obesity. Our current work suggests loss of C2CD5 generates obese phenotype with increased food intake, decreased energy expenditure and an increase in axial length in C2CD5KO (9.2±0.3 vs 10.3±0.2cm) (males; not shown), possibly by the regulation of MC4R trafficking by C2CD5. Injection of melanocortin agonist (MTII) suppressed food intake and MC4R in the PVH are known to be important for this effect [25]. To assess of the role of C2CD5 and MC4R in regulating food intake and energy expenditure, we performed acute, MTII treatment into the PVH (Figure 5). Assessment of parameters of energy balance can be misleading in mice with different BW and composition but body composition does not change during the acute treatment period, effects of BW and composition on energy balance are negligible as each animal acted as its own control.

**Figure 5:**
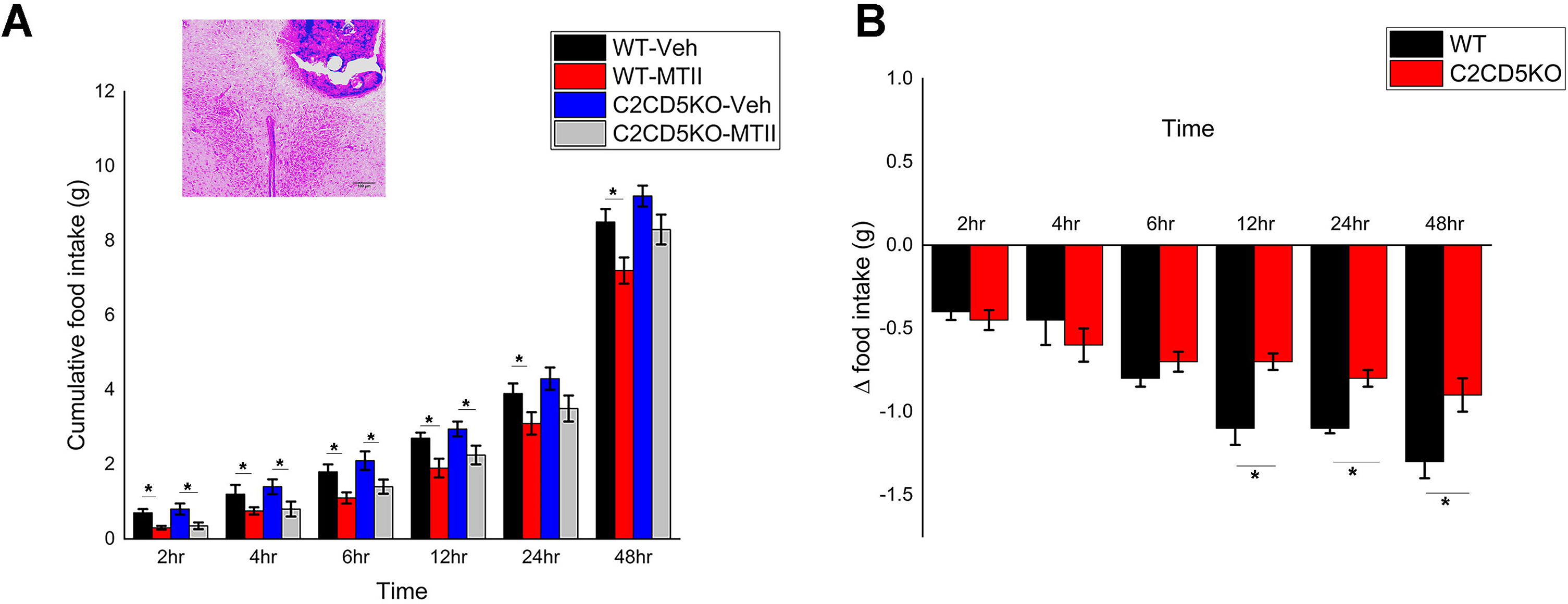
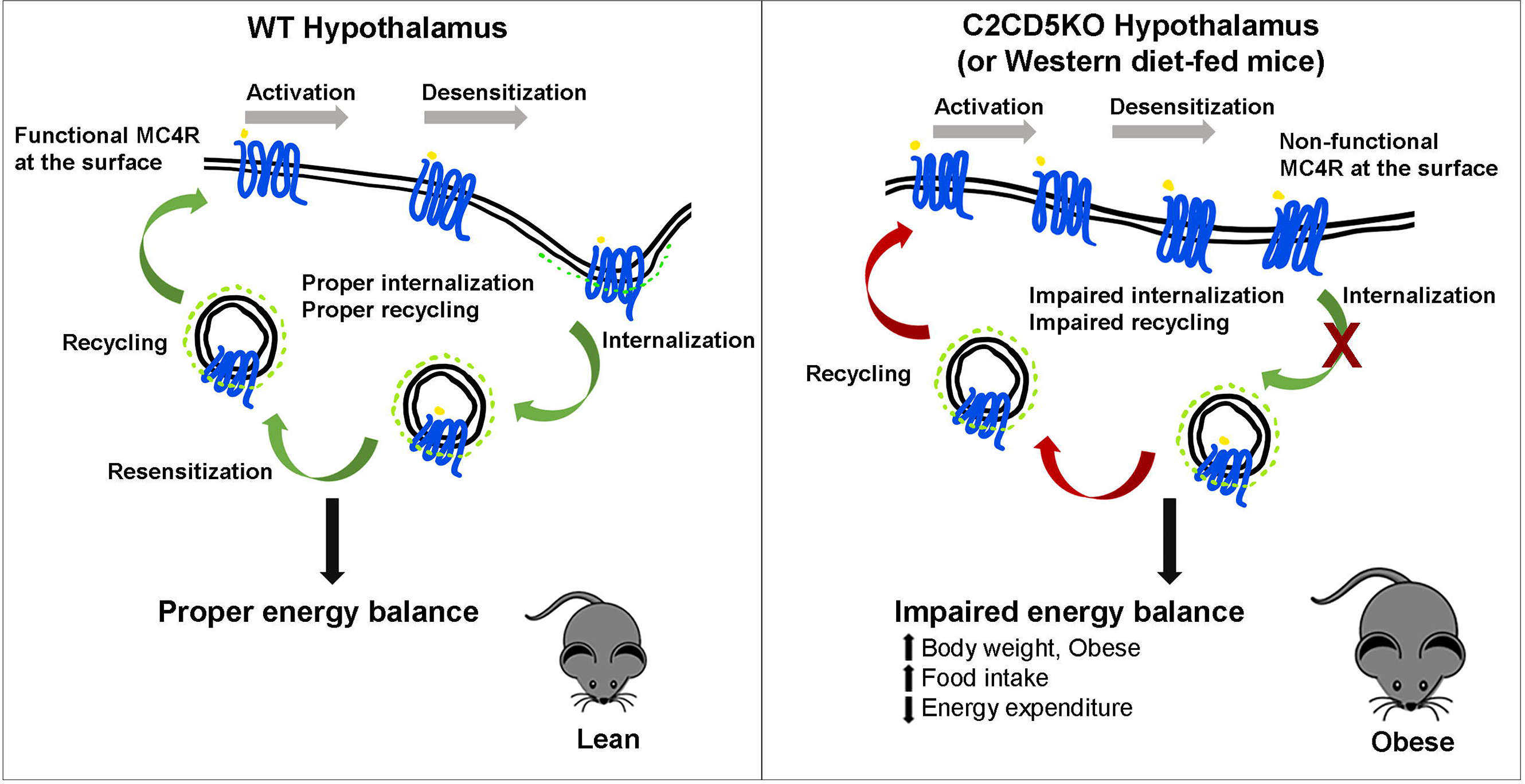
Loss of C2CD5 decreases anorectic effects of intra-PVH MTII. A) Cumulative food intake of WT and C2CD5KO mice after intra-PVH microinjection of vehicle (aCSF) or MTII (n=4/group) (Insert, representative image of intra-PVH cannulation). B) Difference in food intake of WT and C2CD5KO mice treated MTII w.r.t to vehicle. All data are Mean±SEM. *p<0.05

To assess whether the anorectic action of MTII in the PVH neurons is different in C2CD5KO, we injected WT or C2CD5KO male mice with MTII into the PVH and recorded their food intake and energy expenditure for the ensuing 48hrs in metabolic cages. While both WT and C2CD5KO mice reduced their food intake in response to MTII compared to vehicle injected, C2CD5KO mice had a significantly higher food intake at the end of 48hrs irrespective of the treatment (Figure 5A, B). Overall data suggest a role of C2CD5 in the PVH MC4R expressing neurons in mediating MTII’s food intake-reducing effects. MTII has previously been demonstrated to also raise metabolic rate via MC4R [16]. To analyze whether C2CD5 has any effect on this MTII-induced increase in metabolic rate, we also assessed energy expenditure and oxygen consumption in the above treated mice. WT mice increased their oxygen consumption (not shown), energy expenditure, and, total activity in response to MTII (Supplementary 2F, G). However, C2CD5KO mice also increased these parameters in response to MTII, albeit the increase compared to vehicle injection was higher in WT mice (Supplementary 2F, G).

It has been shown that MC4R-expressing neurons in the PVH are downstream of hypothalamic POMC neurons and mediate inhibition of food intake. Although, the mechanisms remain elusive, high levels of circulating estrogens prevent obesity in female mice, providing evidence for sexual dimorphism in the regulation of feeding behavior and energy balance in rodents [52]. For example, compared to male mice, female mice hypothalamus have more Pomc mRNA, POMC projections, and display higher neural activities [53,54]. Recent work by Xu et al., [55] found that loss of the MC4R did not influence estrogenic regulation on food intake, energy expenditure and bodyweight. This may explain the difference in body weight, food intake and energy expenditure between male and female C2CD5KO mice compared to their WT counterparts (Figure 2; Supplementary 1, 2).

The weakness of this study is that we observe that C2CD5 is widely expressed in the brain and presumably regulates multiple central nervous system pathways independent of MC4R. To directly test the role of C2CD5 in the central control of feeding and energy balance, future work using Cre-Lox technology will investigate the consequence of knocking-out C2CD5 from hypothalamic neurons involved in energy balance regulation (such as Sim1-, MC4R-, POMC/NPY-expressing neuronal network) and metabolic tissue (e.g adipose).

The strength of our study is that we demonstrated for the first time that C2CD5 is a trafficking protein nutritionally regulated in the hypothalamus and the expression of C2CD5 is decreased in diet-induced obesity. WD-fed male and female C2CD5KO mice exhibit pronounced obesity partially due to an increase in food intake. A decreased in energy expenditure was observed in male mice only. This Difference in the metabolic phenotype observed in male versus female mice lacking C2CD5, suggest that C2CD5 may be involved in sexual dimorphism observed in metabolic diseases and its treatment efficiency.

C2cd5 and Mc4r are co-expressed in the PVH. MC4R trafficking had been shown to be involved in Obesity and notably, loss of functional C2CD5 impaired MC4R endocytosis and favored the presence of non-functional MC4R at the cell surface. C2CD5KO mice exhibit decreased acute responses to MTII injection into the PVH, suggesting loss of C2CD5 in PVH neural circuitry affecting feeding behaviors. This study suggests that a functional C2CD5 is needed in MC4R expressing neurons for normal regulation of energy balance. A decrease in C2CD5 expression or function would lead to Obesity. In addition, MC4R is the focus of many anti-obesity treatments such as setmelanotide that had been shown to drop hunger in Obese participants. In this study, we identify a new MC4R trafficking protein that could be targeted in Obesity and/or that could mediate some of the effects of anti-obesity drugs known to decrease food intake. It will be of importance to evaluate in future studies the potential of C2CD5 as anti-obesity target in rodent and humans.

## ABBREVIATIONS

MC4R: Melanocortin Receptor 4
C2CD5: C2 Calcium Dependent Domain Containing 5
HA: Hemagglutinin
GFP: Green fluorescent protein
N2A: Neuro2A
α-MSH: α-Melanocyte-stimulating hormone
WT: wildtype
C2CD5KO: C2CD5 whole body knockout
NC: Normal chow
WD: western diet (high-fat/high-sucrose/cholesterol)
MTII: Melanotan II
PVH: Paraventricular hypothalamus
i.p.: intraperitoneal
PBS: phosphate buffered saline
SDS: sodium dodecyl sulfate
BW: body weight
AP2: adaptor protein 2
VO_2_: Oxygen consumption
VCO_2_: carbon dioxide production
RER: respiratory exchange ratio
IP: immunoprecipitation
MS: mass spectrometry
PVDF: polyvinylidene difluoride
TIRF: total internal reflection fluorescence microscope
ACSF: artificial cerebrospinal fluid
FBS: fetal bovine serum
PB: phosphate buffer

## Acknowledgments

We thank Loyola University Chicago animal facility for mice housing. We thank Dr. Giulia Baldini for providing us with N2A (HA-MC4R-GFP) stable cell line. We acknowledge the service of Midwest proteome center, Rosalind Franklin University of Medicine and Science. We would like to thank Dr. Seth Robia for his help with TIRF experiments.

## Author contributions

CKG, Conception and design, Acquisition of data, Analysis and interpretation of data, Drafting, revising, and approved the article. TMC, Acquisition of data, Analysis and interpretation of data, revising, and approved the article. DJR, Acquisition of data, Analysis and interpretation of data, revising, and approved the article. VM-A, Conception and design, Analysis and interpretation of data, Drafting, revising, and approved the article.

## Ethics

This study was performed in strict accordance with the recommendations in the Guide for the Care and Use of Laboratory Animals of the National Institutes of Health. All of the animals were handled according to approved institutional animal care and use committee (IACUC) protocols of Loyola University Chicago. Every effort was made to minimize suffering.

**Supplementary 1:** In female mice, loss of whole body C2CD5 results in obesity upon feeding WD. A) Body weight of female WT and C2CD5KO mice fed NC and WD. Mice were fed WD starting week10 of age just before body weights start to diverge (n=8/group). B) Intra-peritoneal glucose tolerance test in female WT and C2CD5KO after 10 weeks of diet change (n=8/group). C) Intra-peritoneal insulin tolerance test in female WT and C2CD5KO after 10 weeks of diet change (n=8/group). D) Serum insulin levels of female WT and C2CD5KO after 12 weeks of diet change respectively (n=6/group). E) Serum leptin levels of female WT and C2CD5KO after 12 weeks of diet change respectively (n=7/group). F, G) Serum cholesterol and triglycerides levels of female WT and C2CD5KO after 12 weeks of diet change respectively (n=8/group). H) Cumulative food intake in kcal of female WT and C2CD5KO mice before BW diversion when fed NC and WD respectively (n=8/group). All data are Mean±SEM. *p<0.05

**Supplementary 2:** Loss of C2CD5 decreases energy expenditure in male mice but not in females. A, B) Energy expenditure of male and female WT and C2CD5KO mice before BW diversion on NC and WD respectively (n=8/group). C, D) Total day/night activity of male and female WT and C2CD5KO mice before BW diversion on NC and WD respectively (n=8/group). E) Energy expenditure of male WT and C2CD5KO during pair-feeding paradigm (n=4/group). F) Energy expenditure of male WT and C2CD5KO mice after intra-PVH microinjection of vehicle and MTII (n=4/group). G) Total day/night activity of male WT and C2CD5KO mice after intra-PVH microinjection of vehicle and MTII (n=4/group). All data are Mean±SEM. *p<0.05

**Supplementary 3:** A) Agarose gel image showing genotype of C2CD5KO mice (Het, Heterozygous; Hom, Homozygous, KO; Wt, wild-type). B) Western blot image validating C2CD5 antibody using WT, Het, and Hom (KO) mice hypothalamus. C) Electro micrographs showing the validity of immunogold procedure, left – staining in primary omitted hypothalamic sections; right, immunogold staining in C2CD5KO hypothalamic sections; scale 200nm. D) Image showing the pathway analysis of proteins that co-precipitated with C2CD5 in the mouse hypothalamus. E) STRING image showing a global view of proteins and their functional interactions using C2CD5 interacting proteins in mouse hypothalamus.

