## Supplementary figures and images for "Hypothalamic C2-domain protein involved in MC4R trafficking and control of energy balance"

### Supplemental figure1

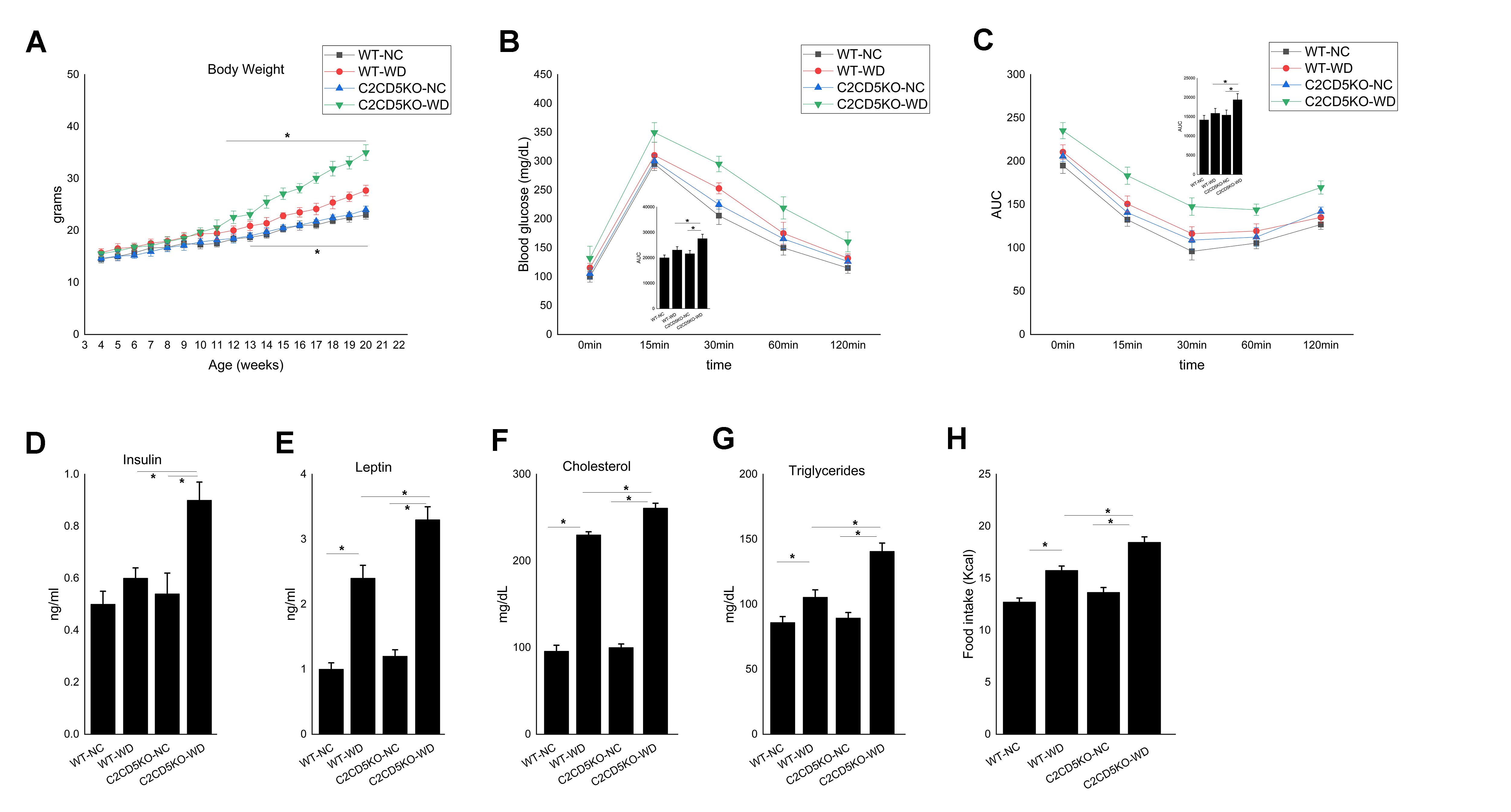

### Supplemental figure2

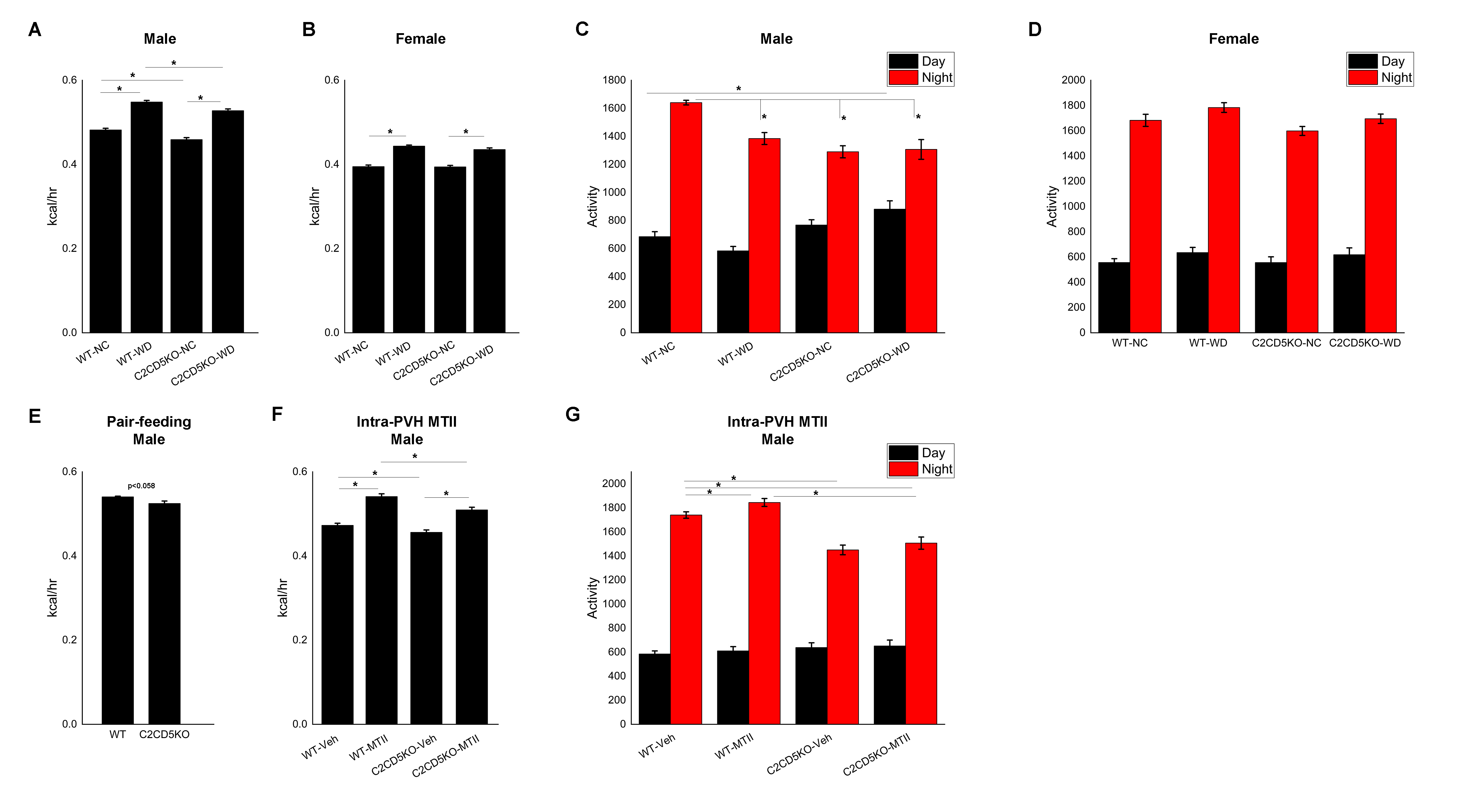

### Supplemental figure3

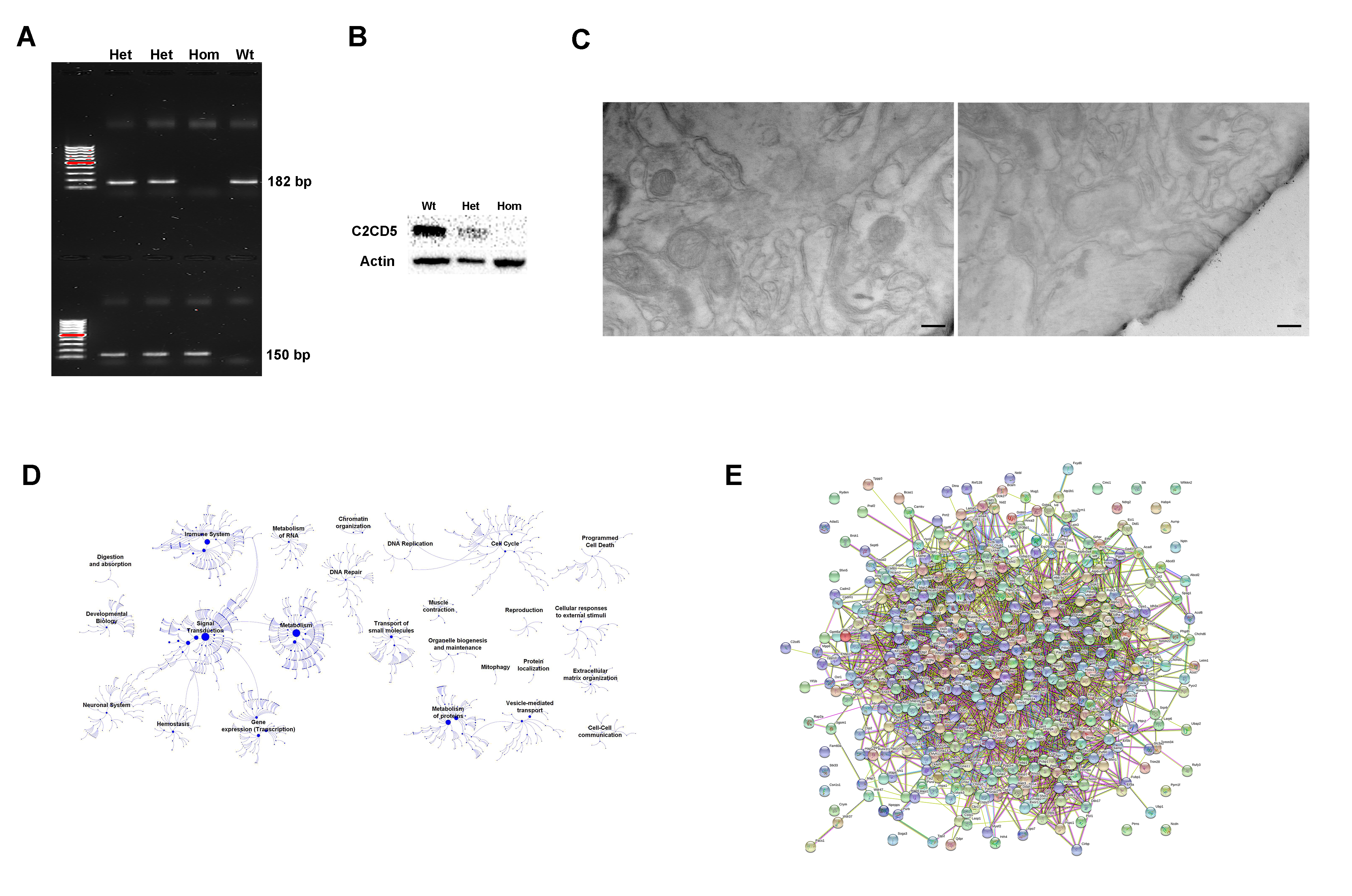
